# Personalized Closed-Loop Brain Stimulation for Effective Neurointervention Across Participants

**DOI:** 10.1101/2021.03.18.436018

**Authors:** Nienke E.R. van Bueren, Thomas L. Reed, Vu Nguyen, James G. Sheffield, Sanne H.G. van der Ven, Michael A. Osborne, Evelyn H. Kroesbergen, Roi Cohen Kadosh

**Author notes:** Shared first author (Nienke E.R. van Bueren, Thomas L. Reed). Corresponding author: Roi Cohen Kadosh.

## Abstract

Accumulating evidence from human-based research has highlighted that the prevalent one-size-fits-all approach for neural and behavioral interventions is inefficient. This approach can benefit one individual, but be ineffective or even detrimental for another. Studying the efficacy of the large range of different parameters for different individuals is costly, time-consuming and requires a large sample size that makes such research impractical and hinders effective interventions. Here an active machine learning technique is presented across participants—personalized Bayesian optimization (pBO)—that searches available parameter combinations to optimize an intervention as a function of an individual’s ability. This novel technique was utilized to identify transcranial alternating current stimulation frequency and current strength combinations most likely to improve arithmetic performance, based on a subject’s baseline arithmetic abilities. The pBO was performed across all subjects tested, building a model of subject performance, capable of recommending parameters for future subjects based on their baseline arithmetic ability. pBO successfully searches, learns, and recommends parameters for an effective neurointervention as supported by behavioral, stimulation, and neural data. The application of pBO in human-based research opens up new avenues for personalized and more effective interventions, as well as discoveries of protocols for treatment and translation to other clinical and non-clinical domains.

## 1. Introduction

There is no doubt that the human organism is complex, and the impact of nature and nurture, as well as their interaction, increases variability between humans. It is therefore not surprising that interventions aimed at altering human behavior are not effective for all individuals. This variability in effectiveness is partly due to the one-size-fits-all approach that currently dominates behavioral intervention research. Accumulating evidence has indicated that this approach is inefficient, and that a treatment that benefits one individual can be ineffective or even detrimental for another individual (1–8). Personalized medicine aims to address this challenge by adjusting treatments to the individual or to a subset of patients (9, 10). Due to the complexity of individual differences, there is an increasing need for personalized medicine for a wide range of drugs, biomedical treatments, and diseases. Without this, the one-size-fits-all approach often only alleviates symptoms in clinical studies without curing the disease (11). This demand for personalization is especially true in the field of transcranial stimulation, where electrical currents targeting specific brain regions are used to alter behavior. Whilst tailoring a stimulation protocol is ideal, identifying the optimal stimulation protocol for an individual proves problematic in large parameter spaces, where the systematic testing of each parameter combination can lead to overly costly and time-consuming protocols. For instance, one stimulation technique that is gaining popularity is transcranial alternating current stimulation (tACS) (12). tACS utilizes an alternating current delivered via multiple electrodes placed on the scalp, which is capable of propagating through the scalp and modulating the activity of the underlying neurons. The applied alternating current promotes oscillatory activity at the stimulation frequency (13), allowing direct modulation of brain oscillations that subserve cognitive processes (14). Through this process, tACS provides an attractive way to investigate causal predictors of behavior and to use such knowledge to improve human capabilities or health. However, exploring the effects of all tACS parameters on the performance of different individuals requires an exhausting amount of testing when considering different current (0-2 mA) and frequency (0-100 Hz) combinations.

One recently proposed method for selecting parameters in brain stimulation is Bayesian optimization (BO) (15, 16). BO is an active machine learning technique that aims to find the global optimum of a black-box function *f*(*x*) by making a series of evaluations. To select the next evaluation, BO first constructs a probabilistic model (surrogate model) for *f*(*x*) and exploits this model to make decisions. This results in a procedure that can find the maximal value of difficult non-concave functions with relatively few evaluations, at the cost of performing more computations to determine the next point (at minimal cost when compared with the effort of evaluating the function at more points) (17). Hence, BO is particularly valuable when there is a need to explore a large experimental space in as few evaluations as possible. Generally, BO involves two procedures: 1) fitting an appropriate model to function *f*(*x*) and 2) choosing an acquisition function α(x) that steers sampling in the direction where improvement over the current best evaluation is most likely.

The present work was inspired by previous work that used BO in human-based research (15, 16, 18–20) In these previous BO studies, all iterations of the process are run on the same individual, allowing the experimenters to achieve person-specific results. However, to do this, the entire BO process must be run for each individual that requires stimulation - a lengthy and costly process. A workable solution is to base the algorithm on a measurable characteristic that varies across subjects, such as baseline ability in the behavioral task of interest. Therefore, we developed a novel personalized (p)BO for human-based research. In pBO, the algorithm is trained on an initial small set of data (burn-in phase), and then iteratively selects stimulation parameters across subsequent subjects, with the aim of identifying the optimal stimulation parameters for improving behavioral performance, whilst considering personalized information. i.e. baseline arithmetic ability (**Figure 1**).

**Figure 1.**
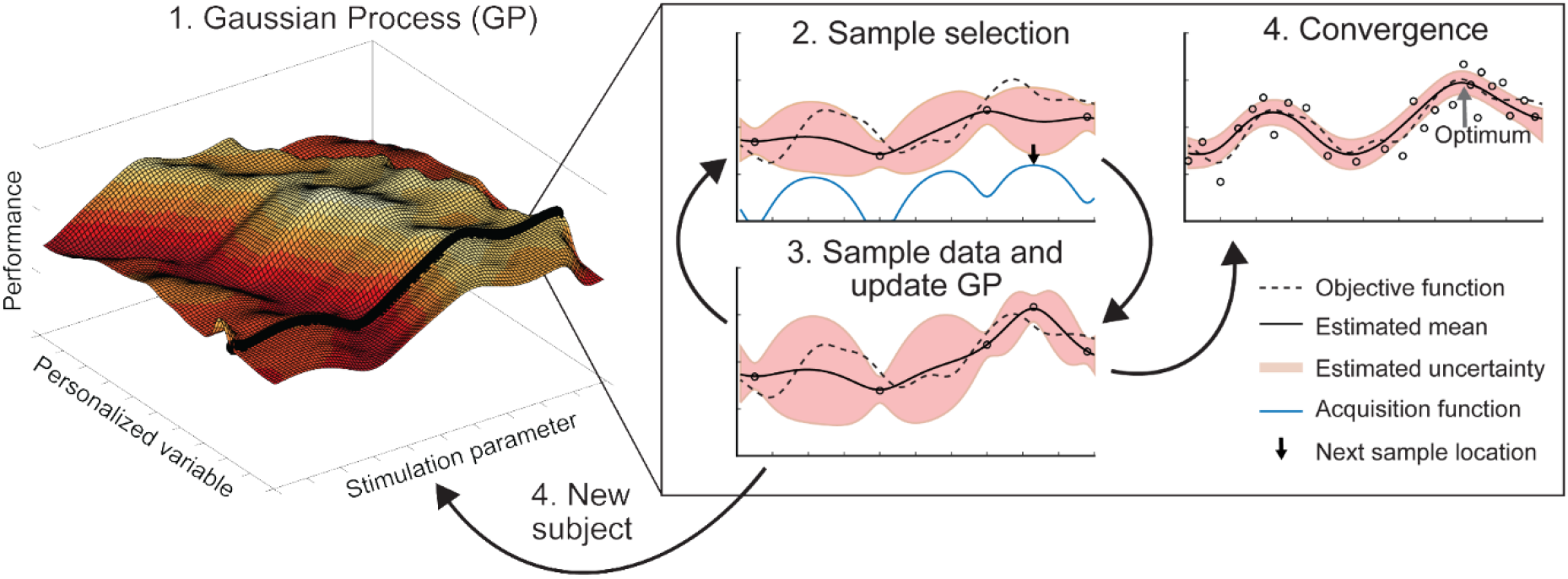
Illustration of the personalized Bayesian optimization procedure**. (1)** The Gaussian process (GP) is fitted to the existing data and models the expected performance along parameter and personalized dimensions. **(2)** The acquisition function identifies the next point to evaluate along the value of the personalized variable relevant to the participant. **(3)** Once the data is collected at this new point the GP is updated and a new point selected. **(4)** This cycle continues until either a new subject is tested, in which case a different value for the personalized variable will be recorded; a pre-set stopping criterion is reached, such as the number of subjects to be tested; or until the potential improvement is considered negligible (convergence). In this study, we utilized a pre-set stopping criterion of 50 subjects, after which testing was ceased.

This BO algorithm can incorporate personalized information (21), including an individual’s data, such as age, gender, neural activity or cognitive profile. Based on the vast literature that highlights the impact of individual differences on stimulation efficacy (22, 23), we personalized the BO to subject’s cognitive ability in this study. We focused on optimizing arithmetic performance considering its importance in the success of one’s future career and socio-economic status (24) and its impairment in acquired and congenital brain disorders (25, 26). Skills needed for solving arithmetic problems vary greatly in the typical and atypical populations (27, 28). Similarly, a recent study on arithmetic skills highlighted the individual differences in both neural correlates and behavioral response in healthy people (29). The left frontoparietal network has been implicated in playing an important role in arithmetic processing and can be targeted by tACS (30, 31). We recognize that other brain stimulation techniques have been used in the field of arithmetic (32–37, for reviews see 31, 38, 39). However, we utilized tACS since this method allows for stimulation at a range of specific frequencies to explore those that might impact arithmetic performance.

We examined whether we could tailor tACS parameters to improve arithmetic performance using a pBO that takes baseline ability into account in healthy subjects. To do this, the individual’s baseline arithmetic ability was initially measured, after which the stimulation parameters to be used were automatically selected either at random, if subjects were in the initial burn-in phase, or by the pBO algorithm. Subjects then completed a block of the arithmetic behavioral task whilst receiving stimulation using the selected parameters (**Figure 2**). Stimulation parameter selection and behavioral testing were repeated in each subject until three blocks of different arithmetic problems were completed. Note that these three blocks were included to select more samples to allow optimization based on the pBO across subjects, rather than optimizing performance over these three iterations. The tACS parameters that were altered were current intensity and frequency, and the pBO algorithm aimed to identify the optimal parameter combination for improving arithmetic ability given a subject’s baseline arithmetic ability. To target the ability to solve arithmetic problems more precisely we used diffusion modeling, which allowed us to target specifically the drift rate, a measure of cognitive ability, rather than auxiliary components such as non-decision response time or response conservativeness (40). Furthermore, we ran different computational simulations to demonstrate the efficiency of our proposed pBO in comparison to random sampling and a standard BO algorithm (i.e., pBO without a personalized variable). We also recorded electrophysiological frequency band power and connectivity at baseline and after applying combinations of tACS parameters to link behavioral changes to electrophysiological (EEG) outcomes, while we report this finding, we note that it is not the main focus of the present study.

**Figure 2.**
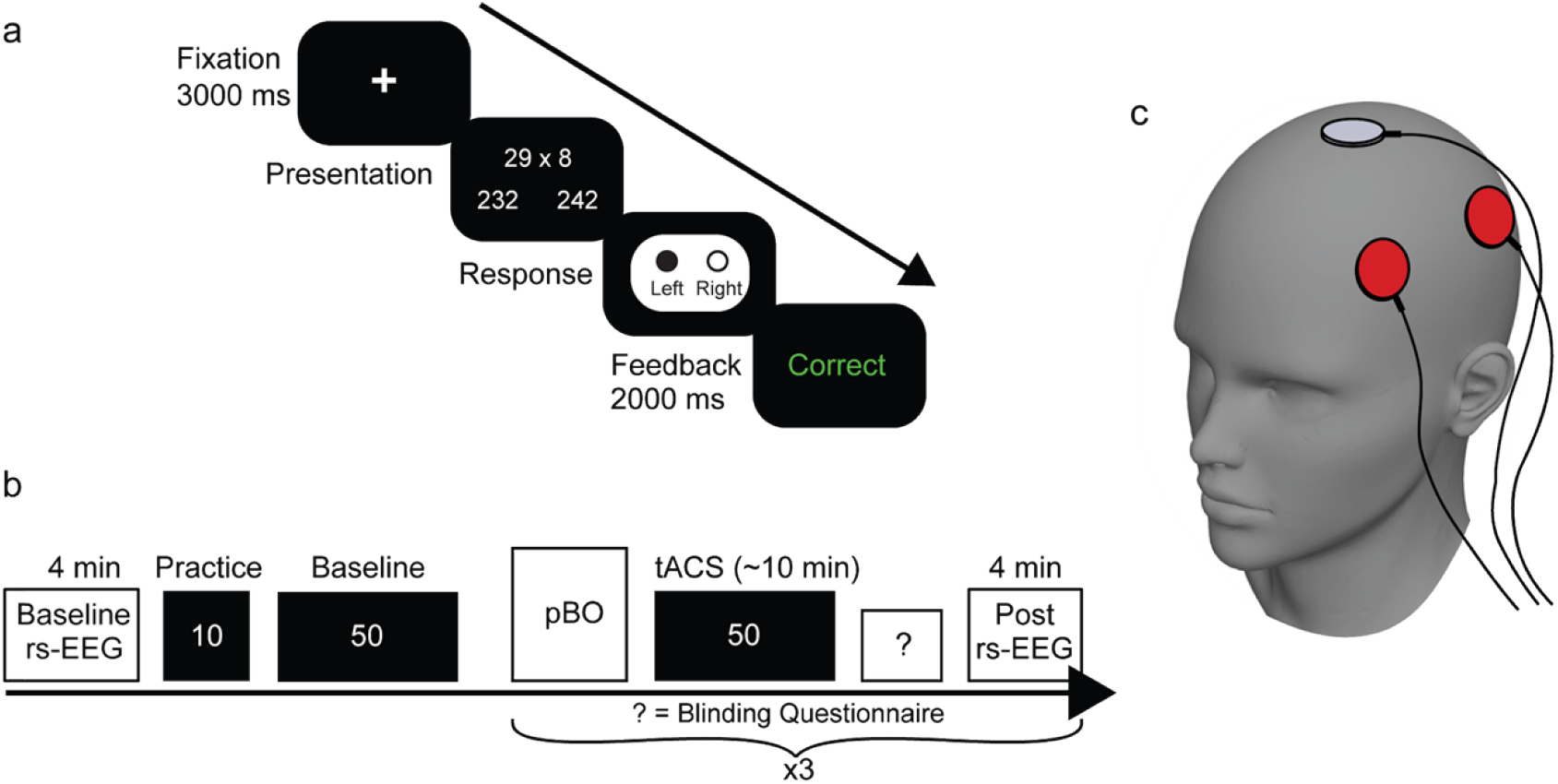
An overview of the experimental paradigm. **a)** An overview of the behavioral paradigm. Subjects (n = 50) watched a fixation point that indicated the start of a trial. After 3000 ms an arithmetic multiplication was shown with two possible answer options on the left and right side with a difference of 10. Subjects responded by pressing either the left or right button on a response box. Lastly, subjects received either ‘correct’ or ‘incorrect’ as feedback to continuously capture attention. **b)** Subjects first completed a baseline rs-EEG of four minutes, after which 10 practice trials of multi-digit times single-digit multiplications were presented. This was followed by the baseline task, which comprised five blocks of 10 different multiplications. Three different tACS frequency-current combinations were proposed by the BO algorithm after the completion of the multiplications. Between these tACS combinations, post-block rs-EEGs were recorded before the subjects moved on to the next tACS combination. Validation of the blinding of the stimulation and perceived sensations were assessed after completion of a stimulation block. **c)** An illustration of the tACS electrode montage. Stimulation was applied over the left frontoparietal area (F3 and P3) with one return electrode (Cz).

## 2. Results

### 2.1. Group-level Personalized Bayesian Optimization

As **Figure 3** shows, the optimal stimulation parameters depended on the participants’ baseline ability: we found a shift from higher frequencies and currents in lower (poor) baseline abilities, to lower frequencies and currents in higher (better) baseline abilities (mean - 1SD (38.67 Hz, 0.97 mA), mean (16.67 Hz, 0.88 mA), mean + 1SD (18 Hz, 0.6 mA), mA values are peak-to-peak). Baseline ability is a continuous parameter and it should be considered that the best inferred tACS combination differs along this continuum.

**Figure 3.**
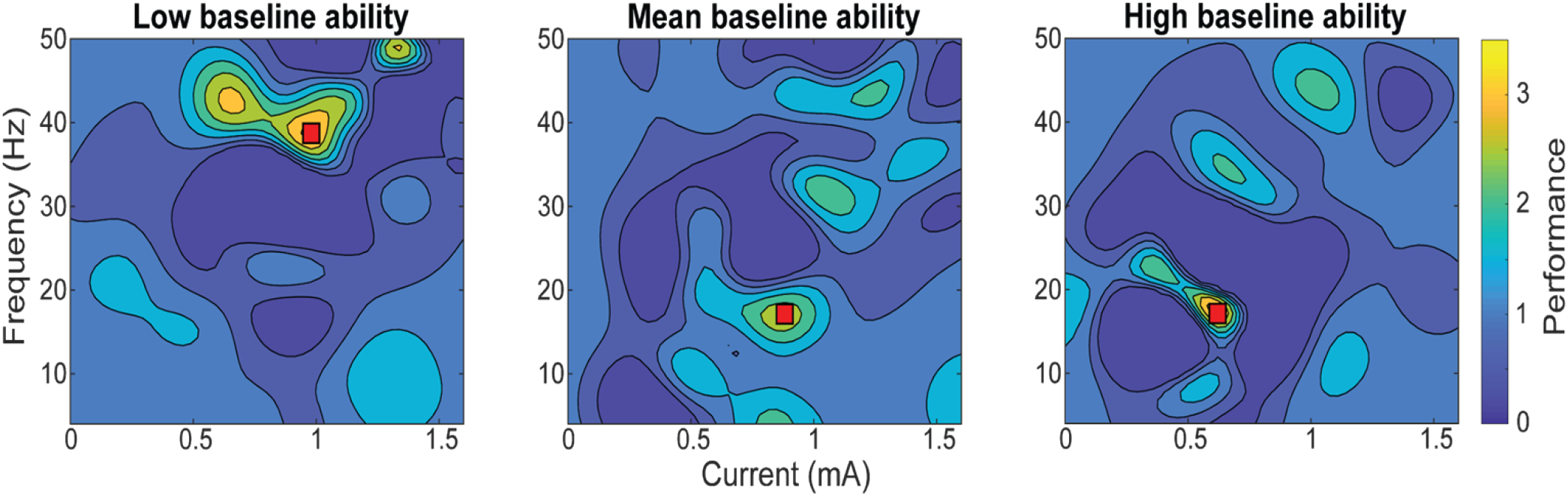
Results of the personalized Bayesian optimization model at several different baseline abilities (n = 49). From left to right, the figure shows the predictions from the Gaussian process model for low baseline ability (1 standard deviation (SD) below the mean), mean baseline ability (mean = 0.055), and for high baseline ability (1 SD above the mean). The y-axis shows the frequency range of the applied stimulation (0- 50 Hz) and the x-axis the current of the stimulation (0-1.6 mA, peak-to-peak). Arithmetic performance is indicated in color based on the normalized drift rates (tACS block/baseline block). Low drift rates are shown in dark blue and high drift rates in yellow. A best-inferred point for arithmetic performance according to a specific frequency-current combination is indicated by a red square in all three panels. Note that this figure is not based on different groups of participants as in moderation analysis, but represents a three dimensional view of the GP’s surrogate surface at three different points for visualization purposes.

A more in-depth visualization of the efficacy of the pBO procedure revealed that the overall fluctuation in performance improvement (e.g., normalized to the baseline performance without stimulation) across subjects with low and high baseline abilities was similar (**Figure 4a**). This result indicates that the success of our approach is equally effective for people with either low or high arithmetic baseline ability. The optimal frequency-current tACS parameter combinations proposed by the pBO algorithm confirms a shift from higher frequencies and currents in low-baseline ability subjects to lower frequencies and currents when baseline ability increases (**Figure 4b**; see also **Figure 3**). In addition, the black-box function *f*(x) is reliably optimized over the course of the iterations, as shown by an increase in the individual’s ability to solve arithmetic problems (**Figure 4c**).

**Figure 4.**
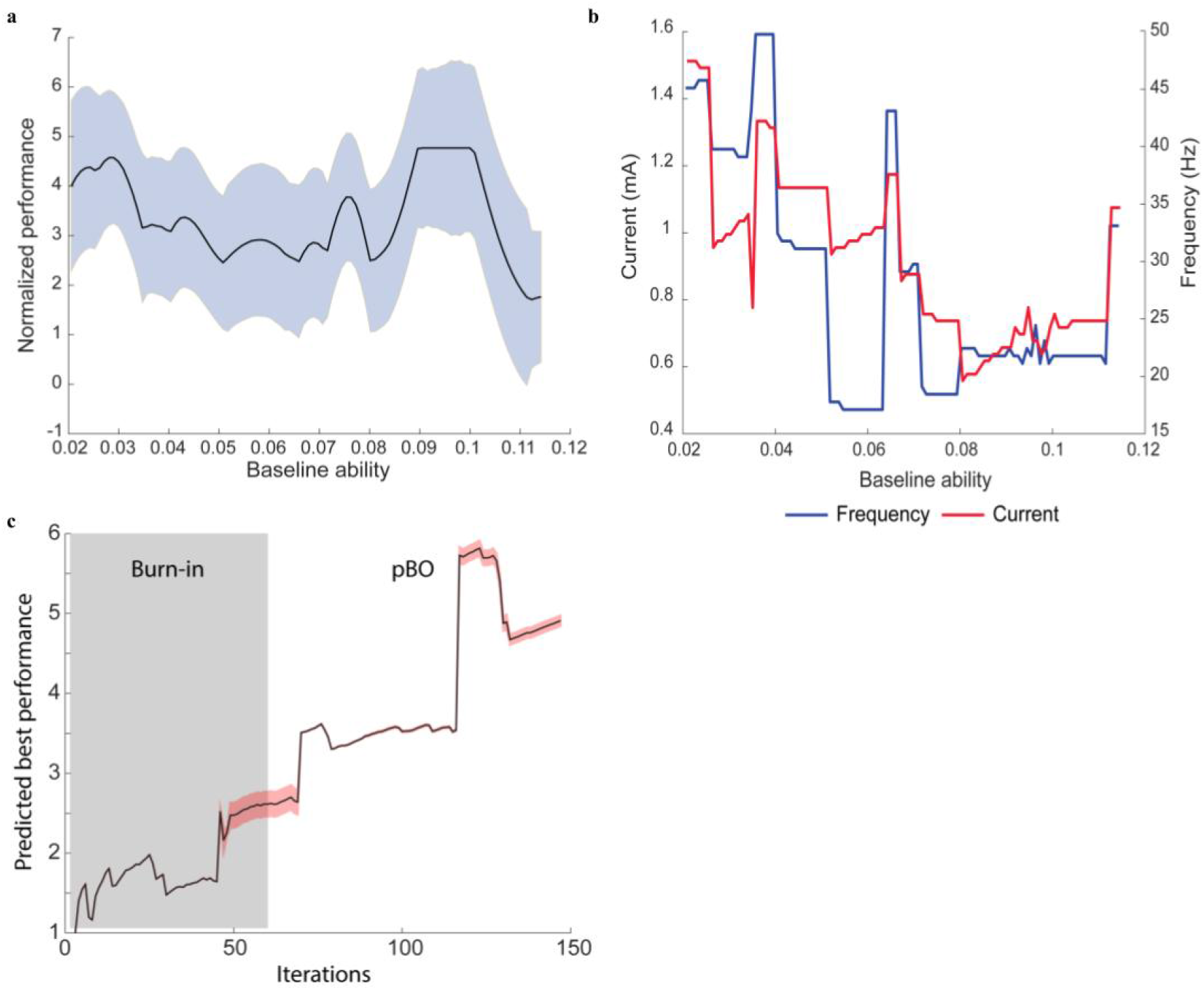
Results of optimizing behavior with personalized Bayesian optimization (pBO) (n = 49). **a)** In-depth visualization of the normalized performance according to baseline ability during pBO. Subjects on the lower part of the baseline ability spectrum showed a similar arithmetic performance improvement during tACS compared to subjects on the higher baseline ability spectrum. Note that a normalized performance score of 1 indicates no difference with baseline arithmetic performance when no stimulation was applied. A normalized performance score higher than 1 indicates improved performance as measured with drift rate. The blue shaded area indicates 95% credibility intervals. **b)** The change in frequency-amplitude tACS parameters proposed by the pBO algorithm based on the individualized baseline ability in arithmetic at the end of optimization. **c)** Predicted best performance at each iteration (i.e., different blocks), calculated as the best performance predicted by the GP at any parameter combination. Three subsequent iterations were assessed for each participant. Surrogate uncertainty is shown by the shaded area in pink. Note that during some iterations uncertainty is higher due to new baseline abilities introduced in the pBO and due to outliers. These outliers are retested later which then reduces uncertainty.

### 2.2. Personalized Bayesian Optimization Simulation Analysis

To demonstrate the efficiency of our proposed pBO, we examined the optimization performance on a Hartmann 3-dimensional function (41). This 3-dimensional function is a suitable benchmark representing our real experiments including three variables (frequency, current, and baseline ability). When running a Hartmann 3-dimensional optimization using the expected improvement (EI) (42) as an acquisition function in the pBO algorithm, the pBO algorithm outperformed a standard BO algorithm as well as random sampling (**Figure 5**). In other words, higher drift rate values are attained more quickly when using the EI pBO procedure in comparison with BO and random sampling (**Figure 5a**), and the pBO algorithm was shown to identify an optima closer to the true optima of the Hartmann function (**Figure 5b**). When the noise variance 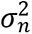 increases, the pBO performance is closer to the performance of random sampling and standard BO (**Figure 5**). As further mentioned in section 4.13 relating to hyperparameter considerations, the estimate of 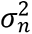 from our observed data which varies by iterations ranged between 0.01 and 2.

**Figure 5.**
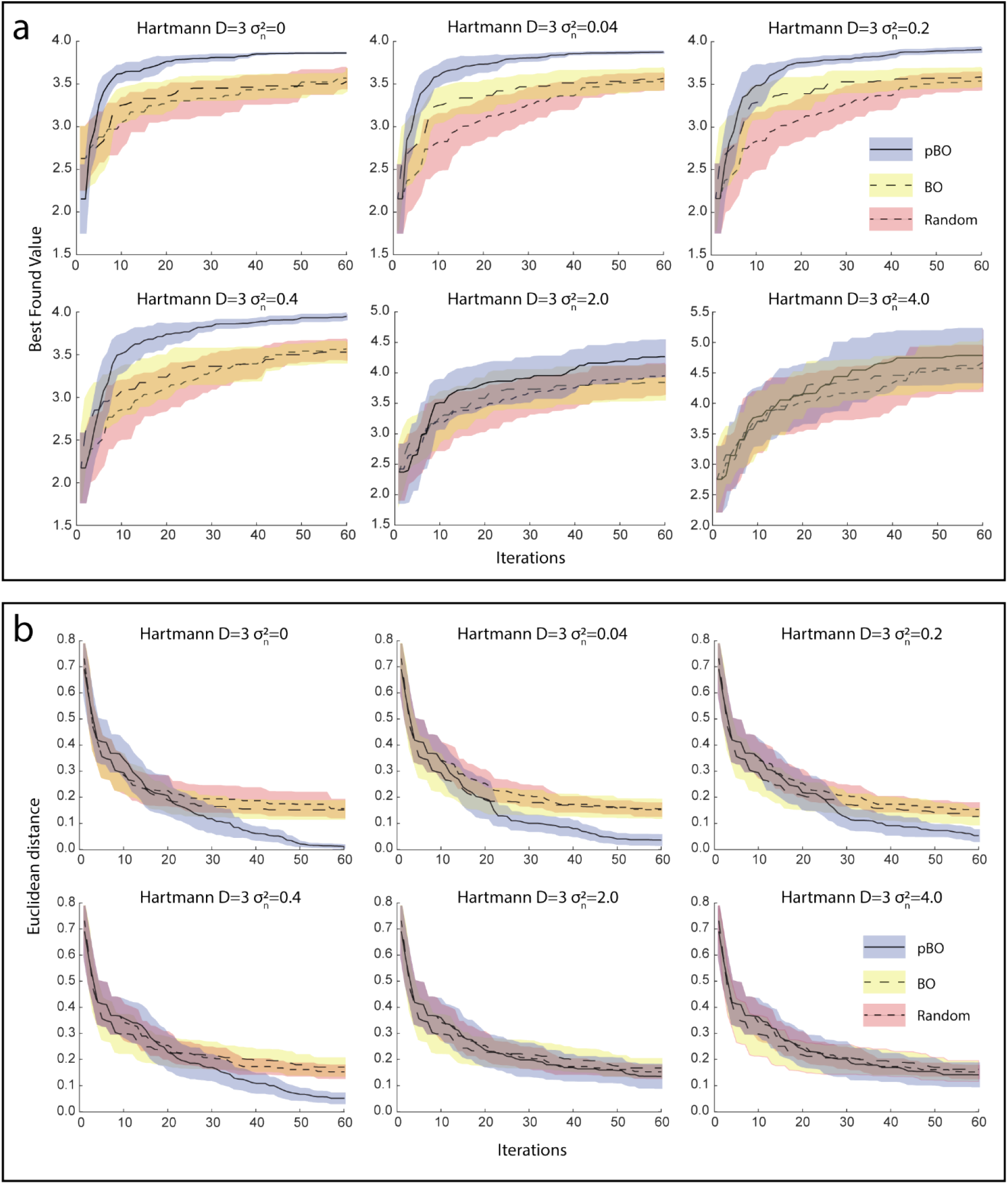
Results of simulating the ability of pBO, BO and random search algorithms on identifying the optima in the Hartmann 3-dimensional surface. The simulation was run 30 times on each of the six different levels of noise, lines represent the mean performance and shaded areas the standard deviation of 30 repeats. **a)** Shows the best found value identified by each algorithm at each iteration, demonstrating that the pBO algorithm is able to find higher values more quickly than the BO and random search algorithms. **b)** Shows the Euclidean distance of the identified optima from the true optima of the Hartmann function (i.e., accuracy of the algorithm). The pBO algorithm is shown to be more accurate than the BO and random search algorithms, except at very high levels of noise, where they are comparable.

Thus, if the behavioral evaluations of the experimental procedure are too noisy, the pBO procedure’s ability to make correct judgements about the optimum parameters is diminished but it is still able to outperform random sampling. Critically, as Figure 5 illustrates within these estimated noise variance ranges our pBO leads to improved optimization compared with the standard BO approach that does not take baseline ability into account. In particular, the standard BO is unable to enhance performance, thus highlighting the benefit of personalization vs. the one-size-fits-all approach.

### 2.3. Baseline Electroencephalography and Arithmetic Ability

To examine whether our results are supported by neurophysiology we used baseline electroencephalography (EEG). Previous EEG studies suggest a positive relationship between left frontoparietal theta (4-8 Hz) connectivity and high-level cognitive processing (43–46). However, none was found when running a regression model trying to predict baseline arithmetic ability from baseline left frontoparietal connectivity in the theta range (all *p* > .3). The same applied to frontal theta power (*p* = .88), theta/beta ratio, and beta (14-30 Hz) connectivity (all *p* > .1). However, our findings from the pBO models highlight an optimal performance effect in the beta frequency range (14-30 Hz) in subjects with average and high baseline ability, whilst low baseline ability individuals benefit from stimulation in the gamma frequency range (> 30 Hz). We therefore examined the relationship between baseline ability and baseline frontal beta power in an exploratory manner. Higher (gamma) frequencies could not be recorded reliably with the equipment. We found that subjects with higher arithmetic baseline ability have higher baseline beta power in comparison to subjects with lower arithmetic baseline skills (non-parametric (Spearman) correlation: *r_s_* = .29, *p* = .03). This neurophysiological finding corroborates the group-level pBO model results, where the pBO algorithm chose the tACS frequency that mirrors baseline neurophysiological activity in those individuals.

## 3. Discussion

Most interventions in humans are not tailored to the individual’s characteristics, such as behavior or brain function, but use a one-size-fits-all approach that leads to inefficient or even ineffective interventions (1–8). This lack of progress is rooted mainly in the complexity of personalizing interventions, due to the immense burden on time and resources. The results of the empirical and simulation experiments performed in the present study demonstrate that our pBO algorithm is capable of tailoring different current strengths and frequencies of tACS to an individual’s baseline ability. Specifically, we demonstrated that the optimal stimulation parameters, determined by the pBO algorithm, differ in low and high arithmetic baseline individuals. This suggests that there are either different cognitive processes involved or differing effects of stimulation in these groups. For example, a recent multiplication study indicated possible behavioral constraints of a one-size-fits-all brain stimulation protocol on performers in a certain subgroup of the population (e.g., high performers) (47). Notably, the present results cannot be explained by a placebo (sham) stimulation, as we controlled for placebo effects of stimulation by including 0 mA in the search space of the pBO algorithm, as well as other very weak currents that are assumed to be ineffective. In addition, if our effects could be explained by a placebo effect, it should have led to the parameters that yielded the strongest sensation, which was not the case (**Figure S2 and S3** in Supporting Information). Similarly, our results cannot be attributed to the participants learning effects as improvement in arithmetic performance was based on an improvement across participants, and not within.

Highlighting the significance of our results, the majority of previous brain stimulation studies that aimed to determine stimulation parameters have only tested a small number of different parameters to observe their differential effects (31). However, this approach leaves a large amount of stimulation parameter combinations unexplored, as such exploration is both expensive and time consuming. Whilst a small number of studies have utilized BO, they have focused on running the entire BO paradigm on one individual to find their best stimulation parameters (15, 16). This approach, while providing many advantages, does not allow for the convenient transfer of parameters optimized for one subject to new subjects, and does not allow its usage in contexts that do not permit repeated sampling in the same individual due to ethical constraints, potential side effects, or time pressure. In contrast, our pBO algorithm provides further advancements by including the following novel processes. Firstly, our algorithm receives personalized data, in this study baseline cognitive data from a subject, and suggests the stimulation parameters to test that are conditioned on the baseline data. Due to the ability to recommend personalized stimulation parameters solely based on a baseline measure, our work on a pBO algorithm represents a significant advance in this area. To illustrate, our experimental findings demonstrate that pBO preferentially selects more successful tACS parameters to optimize the interventional outcome, in this case arithmetic performance (**Figure 4**). Secondly, our study shows that non-personalized interventions, as in standard BO, are ineffective due to the inability to optimize performance effectively (**Figure 5**). Our simulations further show that pBO is reliable even when there is a considerable amount of noise present in the models. Previously, most BO applications have been in a noise-free context, in contrast with human-based studies that are prone to noisy evaluations. More precisely, as noise increases, the pBO algorithm is less able to evaluate the stimulation parameters correctly, but still outperforms random sampling and the standard BO algorithms. This furthermore applies to the estimated noise variance ranges that were observed from our data.

In addition, the behavioral and simulation results support our electrophysiological evidence that pBO can provide new protocols for intervention as well as mechanistic insights. In the present study, pBO highlighted the importance of frontal beta frequencies (14-30 Hz) as indicated by the BO group-level model in subjects with average and high arithmetic abilities. In line with our group-level pBO models, we showed that high baseline ability subjects have higher frontal beta power in comparison with low baseline ability subjects. In addition, subjects with low baseline abilities benefit more from tACS in the gamma frequency range, which is in line with responses linked to spike timing dependent plasticity (STDP) (48). However, gamma frequencies could not be recorded reliably with EEG due to low-pass filtering properties of the skin and skull together with a low signal-to-noise ratio (49), and was therefore not statistically explored. This limitation can be overcome by magnetoencephalography (MEG), as was shown by a study indicating frequency-specific neural entrainment by tACS (50). Our EEG results provide a causal inference of the involvement of baseline oscillatory brain activity due to the overlap between the result of our pBO model and baseline EEG correlates, which in the field of mathematical cognition was so far based on correlation (31, 51).

One constraint of this approach that should be considered, is that the distribution of subject baseline abilities in this study was weighted towards the lower end of the ability range, leading to fewer subjects with higher baseline ability being tested (**Figure S1** in Supporting Information). Therefore, results in the lower baseline ability spectrum are of higher confidence regarding an estimation of the optimum frequency-current tACS combination. This notion is especially relevant on account of the similar results in our group-level pBO model for the mean and high baseline abilities. Additionally, the subjects that participated were mainly university (under)graduates, which might have led to a small arithmetic ability range when compared to the population. A possibility exists that our pBO models will differ slightly when testing more subjects with higher baseline abilities. Moreover, while we demonstrated the personalization of intervention based on two dimensions (i.e., the current and frequency of tACS) and a personalized feature, our algorithm allows for the inclusion of many more dimensions (e.g., phase, brain region, and duration of stimulation) in future interventions. Additionally, the personalized variables are pervasive in human-based research and can include multiple variables such as behavioral data, neural activity, age, and gender. For example, the presented findings could translate to other neurointerventions, such as other forms of transcranial electrical stimulations, transcranial magnetic stimulation (TMS) or to sensing-enabled brain stimulation such as deep brain stimulation (DBS) and the responsive neurostimulation system (RNS). Similar to tACS, these interventions use a broad range of stimulation parameters whereby it is uncertain which parameters are more successful to optimize the interventional outcome due to individual differences in healthy subjects or patients (52–54). Our pBO approach overcomes this limitation by personalizing the intervention based on the selection of stimulation dimensions together with a personalized feature. Taken together, the use of our pBO approach is widely applicable, and can simultaneously model multiple dimensions together with a wide range of choices of personalized variables. Further investigations into closed-loop algorithms for individualized interventions may greatly improve the reliability of those interventions. This is particularly important in a clinical setting where the aim is to optimize symptom improvement.

To conclude, we have demonstrated a more efficient research process, taking as a working model the field of brain stimulation to overcome the problem of selecting stimulation parameters for each individual. The method we suggest here can be extended with minimal or no changes to different fields in which the optimal parameters are unknown and/or expensive to assess, including drug discovery, invasive and non-invasive brain stimulation, and physical and mental training in both typical and atypical populations.

## 4. Methods

### 4.1. Subjects and Ethical Permission

Fifty subjects gave written consent before the start of the study. All met the safety criteria for transcranial electrical stimulation (tES) and received financial compensation of £20. In addition to this compensation, subjects had the chance of winning an additional £50 based on their performance. Behavioral data from all 49 subjects aged between 18-30 years old (31 of whom were female) were used for the pBO (mean age = 22.52 ± standard deviation (SD) = 4.09). All were right-handed. One completed their education at GCSE level, 14 at A-level, 17 were undergraduates and 18 were postgraduates. In the UK educational system GCSE refers to secondary education and A-level refers to an advanced level that can lead to university. All subjects reported no counterindications to electrical stimulation or any history of dyscalculia, dyslexia or attentional deficits. The proposed study received ethical approval from The University of Oxford Medical Sciences Interdivisional Research Ethics Committee (protocol number: MSD-IDREC-C2-2014-033). Additionally, we pre-registered the present study on the Open Science Framework; see https://osf.io/bg2pd.

### 4.2. Overview of Experimental Paradigm and Stimuli

Over the course of the experiment, subjects completed four blocks of fifty multiplication problems - one baseline block and three stimulation blocks (see **Figure 2**). After recording an initial 4-minute baseline resting state EEG, the task was explained to the subjects and they completed 10 practice trials, followed by the baseline block during which no tACS was administered. In each block, subjects had to indicate which answer was correct as accurately and as fast as possible (see **Figure 2a**). Subjects indicated the correct answer by pressing either the left or the right button on a response box situated in front of them. They underwent three blocks of multiplications in which they received tACS. Prior to each tACS block, the pBO algorithm was run to determine the stimulation parameters (current intensity and frequency) to be delivered during the upcoming experimental block based on the individual subject’s performance in the baseline block. The stimulation parameters were automatically selected by the algorithm and administered whilst maintaining blinding in both the subject and experimenter (for a complete list of the applied current and frequency, see **Tables S1 and S2**, in the Supporting Information. Rs-EEG was recorded again after each stimulation block.

### 4.3. Behavioral Stimuli

Arithmetic performance was tested using an arithmetic calculation paradigm, consisting of problems involving a single-digit number multiplied by a two-digit number, with a three-digit outcome. A calculation paradigm was used instead of a retrieval paradigm since calculation has been associated with an increased activation in the frontoparietal network (30, 55, 56). None of the multiplications included operands with the digits 0, 1, or 2 to prevent variations in difficulty. In addition, the two-digit operand was not smaller than 15, did not use repeated digits, and was not a multiple of 10. Subjects were visually presented with a multiplication problem on a screen with a correct and incorrect answer positioned under the multiplication problem on the left and right side. The position of the correct and incorrect answer was randomly allocated to the right and left sides of the screen and they always differed by 10. Each problem was presented only once, and their order was randomized.

### 4.4. Measurement of Baseline Abilities

An arithmetic baseline task containing 50 different arithmetic multiplications was presented to measure individual arithmetic ability in terms of response times and accuracy. Subsequently, baseline drift rates were calculated for each subject according to the two-choice EZ-diffusion model (40). This approach allowed us to dissect the different components in the chain of information processing by modeling the decision process and targeting the cognitive component of interest (the drift rate, which reflects ability and task difficulty by modeling response time and accuracy), rather than auxiliary components (40, 57). This model was chosen to combine reliably the response time and accuracy in one outcome that could be optimized through the pBO procedure. The 50 trials completed in the baseline block were randomly divided into two sets, and for both sets a separate drift rate was calculated. One was used as a measure of the subject’s baseline ability, whilst the other was used to normalize the drift rates calculated during the optimization phase (e.g., during the experimental procedure of the pBO). This was done to eliminate dependency between the subject’s baseline ability score and the normalized score in each stimulation block (58). To reduce fatigue, subjects had a break of 30 s after every 10 trials. After completing the baseline task, subjects had a short break (∼3 minutes) before they continued with the experimental procedure of the pBO.

### 4.5. Experimental Procedure of the pBO

Before the start of the pBO procedure, a burn-in phase was used that consisted of 60 random tACS parameters assigned to the first 20 subjects to initiate the pBO procedure. After the burn-in phase, the pBO procedure was run prior to each behavioral block subjects completed. The pBO algorithm selected stimulation parameters for the subject, with the aim of improving behavioral performance given their baseline ability. This was done in an iterative process across 30 subjects, with the algorithm’s estimate of the optimal stimulation parameter, at any given baseline ability, becoming more accurate as more subjects were tested. At each run of the pBO algorithm, all previously collected data was used, including data collected in the burn-in phase and the GP was refitted to model all the data. As we a priori defined in our preregistration, we utilized a pre-set stopping criteria of 50 subjects, after which testing was ceased.

Our rationale to set the sample size to 50 subjects was as follows: For BO without noise (59), n=10–20 per dimension is often used. For BO with noise, a recent work set the number of evaluations to 25 per dimension (60). However, to take into account the possibility that we might deal with increased noise in the present study, we set it to n=50 per dimension (50 subjects x 3 blocks each, equals 150 evaluations to account for three dimensions (frequency, current, and baseline ability).

In total, 150 diverse multiplication problems (three blocks of 50 trials) were administered during the experimental procedure. After each block, performance drift rates were calculated immediately, another rs-EEG was measured for four minutes, and then for the next block the combination of tACS parameters (frequency and current) was changed. Thus, behavioral performance optimization relied on the frequency and current of tACS together with the baseline cognitive ability as indicated by the drift rate. Each subject received three different frequency-current tACS combinations.

### 4.6. Transcranial Alternating Current Stimulation

The alternating current stimulation was administered over the left frontoparietal network (see **Figure 2c**). The tACS was delivered via two stimulation (3.14 mm diameter) NG Pistim Ag/AgCl electrodes (F3 and P3) with one return electrode (Cz) using the Starstim 32 (Neuroelectrics, Barcelona). The impedances of the electrodes were held at < 10 kΩ. The stimulation intensity ranged between 0.1-1.6 mA peak-to-peak in steps of 0.1 for the burn-in phase of the study. For the optimization phase, 0 mA was added to control for possible sham influences. We chose the maximum stimulation intensity based on a small pilot study on three subjects to determine the maximum comfortable intensity. The stimulation frequency ranged between 5-50 Hz in steps of 1 Hz for the whole experiment.

Stimulation was administered in a double-blind manner during the three experimental blocks with a maximum of 10 minutes for each block. Stimulation started 45 s before the start of the block and changed after every block. If the subjects received a stimulation intensity of 0 mA during a block (sham stimulation), a ramp-up and a ramp-down of 30 s was initiated to provide the initial skin sensations during stimulation to ensure blinding. When the subject completed a block within 10 minutes, stimulation was ramped down for 30 s and the subject proceeded to the four-minute rs-EEG. Note that in cases where subjects completed the task in under 10 minutes, they did not receive the full length of stimulation. These stimulation blocks did not differ from the stimulation block in which the subject received the full length of stimulation except for performance. Twenty-four of the 150 stimulation blocks had a duration of less than 10 minutes but more than 7.50 minutes, and 126 stimulation blocks had a duration of more than 10 minutes. This posed no problem, since the present study only investigated the online effects of tACS on arithmetic behavior. After completing a block related to one tACS combination, the subjects filled out a questionnaire in which they were asked several questions designed to gauge the level of sensation experienced during stimulation (see the Supporting Information for the full questionnaire). We used this data to assess the relationship between the intensity rating of every sensation and tACS amplitude.

### 4.7. Resting State-EEG Recordings and Pre-processing

Resting state-EEG recordings were made at the start of the study (before baseline measurements) and immediately after every stimulation block. Electrophysiological data were obtained with eight gel Ag/AgCl electrodes (F3, P3, F4, P4, Fz, Cz, Pz, AF8) according to the international 10/10 EEG system using the wireless Starstim R32 sensor system (Neuroelectrics, Barcelona, Spain), with no online filters. The ground consisted of adhesive active common mode sense (CMS) and passive driven right leg (DRL) electrodes which were positioned on the right mastoid. All EEG measurements had a duration of four minutes in which the subjects had their eyes open while watching a fixation point in the middle of the screen. Raw EEG data were recorded and used for offline analysis using EEGLAB 13.6.5b (61), which is an open source toolbox running on Matlab R2018b (64). Data were high-pass filtered at 0.1 Hz, and low-pass filtered at 50 Hz. Visual inspection was carried out to remove artefacts caused by muscle movement. We rejected an EEG recording from analysis if more than 25 percent of the data in a given block were removed. This resulted in four rejected datasets (2% of the data). Independent component analysis was used to remove blinks and noisy components. On average, 1.23 ± 0.62 SD components were rejected per subject, with a maximum of three components and a minimum of zero.

### 4.8. Sensation Analysis

In line with a previous study (62), and as stated in our pre-registration, we expected higher sensitivity ratings of the tACS parameters with higher current values compared with lower currents. Therefore, we predicted a positive correlation between the intensity rating for itching, pain, burning, phosphenes, warmth, and fatigue. We performed a separate correlation analysis to calculate the bivariate Pearson’s coefficient (r) or Spearman’s rho (r_s_) depending on normality to assess the relationship between the intensity of different sensations induced by tACS and height of tACS current (**Figures S2** and **S3** in Supporting Information).

### 4.9. Bayesian Optimization

Bayesian optimization uses *f* to denote an unknown objective function (e.g., black-box function) for which we do not have a closed-form expression, but we could have an infinite number of queries. Furthermore, this black-box function is expensive and time costly to evaluate. Formally, let *f*: *X*→ *R* (*R* is the set of all real numbers, representing the values from −∞ *to* + ∞) be a well-behaved function, defined on a subset *X* ⊆ *R*^*d*^ *whereby d is the number of dimensions*. The standard BO approach is aimed at solving the following global optimization problem:

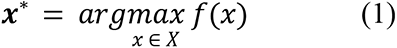

The BO algorithm is aimed at finding the global optimum of arithmetic performance, as indicated by drift rates, of the black-box function *f*(*x*) by making a series of evaluations at *x*_1_,*x*_2_,…,*x*_*T*_.

### 4.10. Personalized Bayesian Optimization

While our approach could personalize a given treatment based on any individual characteristic, such as neural or biometric data, we chose cognitive ability, as the literature provides more supporting evidence for its moderating effect (1–3, 7, 8, 45, 63), especially as it is closely related to our desired behavioral outcome. Each subject has their individual arithmetic baseline ability: this value is considered as the personalized value. We needed to measure this value *p* separately for every subject, as was done during the baseline task. We expected to see that the optimal parameters will vary with different baseline abilities. That is, the optimal parameter *x*^∗^ depends on the different values of *p*. For this reason, the standard BO presented in the previous section may not have been appropriate. Therefore, we proposed to solve the following optimization problem, defined formally as:

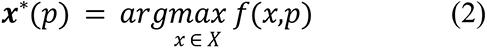

where *p* is the baselines ability given for each subject. The optimal parameter *x*^∗^ is not defined globally, but specifically to a variable *p*. This is the key difference of our pBO in comparison with the standard BO, while we acknowledge related research in BO with environmental variables (21, 65, 67).

### 4.11. Objective Function

In the present tACS study, we used the following objective function *f*(*x*,*p*):

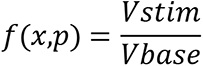

where Vstim is the drift rate during the 50 trials of an arithmetic multiplication block normalized by the Vbase (the drift rate calculated over 25 random trials from the baseline task). To determine improvement between the baseline and the stimulation block, a second Vbase was calculated over the other 25 trials from the baseline task. A Pearson correlation was calculated to determine if the two drift rates from the baseline and the one used for the improvement index are related (*r* = 57, p < .001). The acquisition functions are carefully designed to allow a trade-off between exploration of the search space and the exploitation of current promising regions. A burn-in phase of 60 random tACS frequency-current combinations was used. These were assigned to the first 20 subjects of the BO design to determine the amount of variation induced by stimulation. We decided to use a large burn-in in our paradigm to design a reliable BO algorithm that was based on a large amount of data.

### 4.12. Personalized Gaussian Process for Joint Modeling of Target Function and Baseline Ability

Standard BO models *f* with a GP, *f* ∼ *GP*(*m*,*k*), where *m* is the mean function and *k* is the covariance function (63). This flexible distribution allowed us to associate a normally distributed random variable at every point in the continuous input space. Therefore, we obtained the predictive distribution for ***f*** at a new observation *x* that also follows a Gaussian distribution. Its mean (µ) and variance (*σ*^2^) are given by:

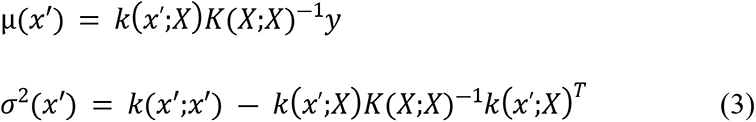

where *K*(*U*;*V*) is a covariance matrix whose element (*i*;*j*) is calculated as *k*_*i*,*j*_ = *k*(*x*_*i*_; *x*_*j*_) with *x*_*i*_ ∈ *U* and *x*_*j*_ ∈ *V*. Behavioral observations are typically associated with noise that can be accommodated in a GP model. Namely, every *f*(*x*) processes extra variance due to independent noise:

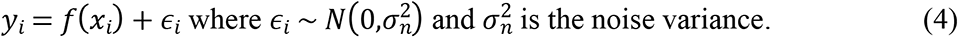

When considering noise, the output follows the GP as 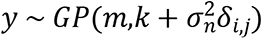, where *δ*_*i*,*j*_ = 1 if *i* = *j* is the Kronecker’s delta. The covariance function for a noisy process becomes the sum of the signal covariance and the noise covariance. Specifically, the exponentiated-quadratic covariance function between two observations can be computed as:

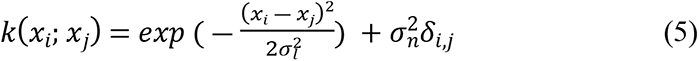

For a more elaborate overview of GPs, we refer the interested reader to Rasmussen and Williams (66).

In our personalized setting, one of the possible solutions is to build a GP and optimization for each value *p*. However, such a simplistic approach faces a critical problem of data efficiency, because the number of data samples is not sufficient to estimate each value *p* separately. Therefore, we extended the GP surrogate to jointly model our target function *f* and the additional personalized dimension *p*, rather than using a separate GP for every subject. Specifically, the GP covariance becomes:

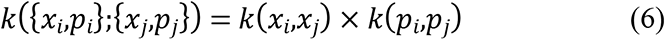

where *k*(*x*_*i*_,*x*_*j*_) is defined in equation (5) and 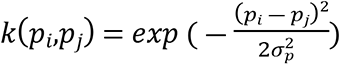. These covariance functions correspond to the parameters and baselines, respectively. We note that the length scale parameter 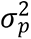 used in *k*(*p*_*i*_,*p*_*j*_) is different from 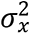 used in *k*(*x*_*i*_,*x*_*j*_). For example, if the baseline ability length-scale 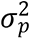 is extremely large, it means the performance is not changing with respect to the baseline performance. On the other hand, if 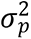 is small, it means the performance function is changing rapidly with the baseline performance. We later maximized the marginal likelihood to estimate these length scale parameters directly from the data (66). Under our modification for the GP, we could estimate the predictive mean and predictive variance:

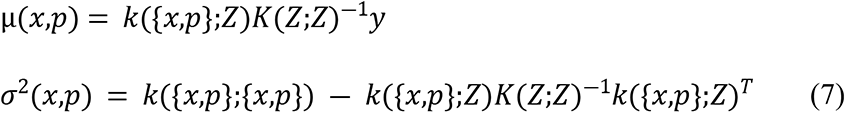

Where we denoted *Z* = [*X*,*P*], the personalized covariance matrix *k* is defined in equation (6).

### 4.13. Acquisition Function

To select the next point to evaluate, the acquisition function *α*(*x*) was chosen to construct a utility function based on the GP surrogate model mentioned above. Instead of maximizing the expensive original function *f*, we maximized the cheaper acquisition function to select the next most optimal point:

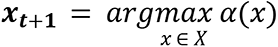

In this auxiliary maximization problem, the acquisition function form is known and can be easily optimized by standard numerical techniques. One of the most common choices for the acquisition function is the GP upper confidence bound (GP-UCB):

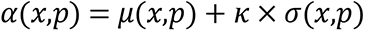

where *μ*(*x*,*p*) and *σ*(*x*,*p*) are the GP predictive mean and variance defined in equation (7) and *κ* is the hyperparameter controlling the exploration-exploitation trade-off. One can follow Srinivas et al. to specify the value of *κ* to achieve the theoretically-guaranteed performance (67). The second common acquisition function is the expected improvement (EI) (42). The EI finds the next sampling point given the highest chance of expectation to improve upon the best-found value so far. Using the analytical expression of Gaussian distribution, we have the EI in closed-form as:

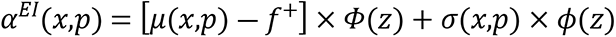

Where 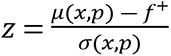 and *f*^+^ is the best observed value up to the current iteration, *Φ*(*z*) is the standard normal cumulative distribution function and φ(*z*) is the standard normal probability density function. When the uncertainty is zero *σ*(*x*,*p*) = 0, the *α*^*EI*^(*x*,*p*) = 0.

#### Hyperparameters considerations

Personalized Bayesian optimization relies on a personalized Gaussian process surrogate model to select a next point for testing. This personalized GP model (defined in Section 4.12) involves several hyperparameters including the length scales *σ*_*l*_ in Eq. (5), *σ*_*p*_ in Eq. (6), and the noise variance 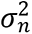 in Eq. (4). We make use of the property of the Gaussian process to estimate these hyperparameters directly from the observed data by maximizing the log marginal likelihood of a GP model (65). For robustness, we have also normalized the input *x* ∈ [0,1]^2^, *p*[0,1] and standardized the output score *N*(0, 1) as popularly used in previous work (67). Given this normalized space, the estimated hyperparameters vary by iterations within the range as follows *σ*_*l*_ ∈ [0.03,0.4], *σ*_*p*_ ∈ [0.07,0.5] and 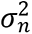 ∈ [0.01,2].

One can optimize the EI (42, 70) over the current best result or the GP-UCB (27). In short, it is more likely that the UCB selects evaluations with both a high mean and high variance. The EI and UCB have been shown to be efficient in the number of function evaluations required to find the global optimum of many multimodal black-box functions (67, 69). During the present study, the EI was applied to find the optimum in arithmetic performance. Lastly, we decided to remove one extreme drift rate value of 3.6 during the experimental procedure due to possible ceiling effects of the BO for sampling the same stimulation parameters. However, the Supporting Information shows that the inclusion of this data point did not significantly alter our results. For similar results without exclusion of this data point, see **Figure S4** in Supporting Information. In total, we acquired 148/150 iterations. In addition, due to technical problems another data point was not included in the BO procedure.

### 4.14. Simulation Analysis

Simulations were run to validate the pBO procedure during arithmetic performance and tACS. This analysis aimed to show that pBO can outperform both a ‘standard’ BO algorithm and random sampling when identifying an optima in a noisy environment. Note that the present study contained three dimensions, namely frequency, current, and individualized baseline ability. Therefore, we utilized a Hartmann function (41) that included four local minima in three dimensions as an example, to enable our simulations to be comparable to our experimental data. As human-based studies are prone to noisy evaluations, we decided to introduce noise in the simulation by running the same Hartmann 3-dimensional function whilst adding different noise variation values 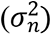. The pBO algorithm presented in this work was compared to a standard BO algorithm which did not incorporate the personalized dimension into its evaluations, as well as a random sampling algorithm. Performance in these simulations was compared in terms of the best found value at each of the 60 iterations, as well as the distance from the known optima location with the Euclidean distance as a metric. Each simulation was repeated 30 times at each level of noise, and the three algorithm’s mean performance and standard deviation over these repeats was calculated. Note that it is not possible to calculate the Euclidean distance between subsequent stimulation pairs due to the inclusion of a personalized variable.

### 4.15. EEG Analysis of Spectral Power and Frontoparietal Theta Connectivity

The rs-EEG data of the remaining datasets were separated in 2 second segments with an overlap of 1 second and windowed with a Hann window. Subsequently, data were transformed into the frequency domain via fast Fourier transformation (FFT). Theta (4-8 Hz) and beta (14-30 Hz) frequency bands were calculated according to their relative power (μV^2^) and normalized by dividing the absolute frequency power of each frequency band by the average absolute power in the 1.5-30 Hz range. In addition, we also normalized the power by dividing the absolute frequency power by the average absolute power in the 4-50 Hz range. The weighted phase lag index (wPLI) in the theta and beta range was computed to determine the phase lag synchronization between the left frontal and parietal areas at baseline and after every tACS block. This computation was made for the complementary channels F3 and P3. The theta wPLI was calculated for 4-8 Hz in steps of 1 Hz and the beta wPLI was calculated from 14-30 Hz in steps of 4 Hz. Furthermore, we normalized wPLI by calculating the wPLI at the applied tACS frequency divided by the baseline wPLI at the same frequency.

First, outliers were removed with Cook’s distance before running statistical models. To focus on the relation between arithmetic baseline ability and spectral power, separate regression models were run with theta and beta power as dependent factors. Likewise, we tested whether there was a relation between frontoparietal theta and beta connectivity scores by running several regression models in steps of 1 Hz for theta wPLI and steps of 4 Hz for beta wPLI.

### 4.16. Statistical Analysis

All the reported inferential statistical analysis was done with RStudio version 1.2.5042 with significance defined as *p* ≤ 0.05 (71). All data is presented as mean ± SD with n = 50 for electrophysiology analysis and n = 49 for the pBO analysis. Pre-processing of electrophysiological data was done with EEGLAB 13.6.5b (61) which is an open source toolbox running on MatlabR2018b (64). Subsequently, normalized electrophysiological data was checked for outliers with cook’s distance and entered in log-transformed regression models with spectral power or connectivity measures as dependent variables and arithmetic baseline ability as independent variable. A correlation analysis on normally distributed datasets was run to calculate the bivariate Pearson’s coefficient (r) to investigate differences in sensation and blinding, and a non-parametric (Spearman’s rho (r_s_)) correlation on non-normalized datasets. Generalized linear mixed effects models (GLMM) were run to explore EEG changes induced by tACS during arithmetic performance (see Supporting Information) The pBO algorithm and simulations were run with Python version 3.6 (72). Note that no inferential statistics such as a GLMM is able to reliably investigate performance gains due to the inability to disentangle the exploration and exploitation trade-off of the pBO algorithm between blocks.

## Acknowledgements

The work for this study was supported by the Prince Bernhard Culture grant for young researchers awarded to NERvB in 2018. The Wellcome Centre for Integrative Neuroimaging is supported by core funding from the Wellcome Trust (203139/Z/16/Z).

## Author Contributions

RCK initiated the idea of the present study; NERvB, TLR, and RCK designed the research; VN provided the code and wrote the associated section in the manuscript; NERvB performed the experiment; NERvB, TLR, VN, JGS analyzed the data or provided help; NERvB, TLR, VN, and RCK wrote the manuscript; and all authors read and approved the final version.

## Data Availability

The datasets generated and analyzed during the present study are available from https://osf.io/bg2pd.

## Code Availability

Matlab, RStudio, and Python code used in this study will be available upon acceptance from https://osf.io/bg2pd and Github.

## Competing Interests Statement

RCK consults, and serves on the scientific advisory boards for Neuroelectrics and Innosphere. The UK Patent Application Number 2000874.4 (“method for obtaining personalized parameters for transcranial stimulation, transcranial system, method of applying transcranial stimulation”) was filed by NERvB, TLR, VN, and RCK. This patent covers the method of obtaining personalized parameters for transcranial stimulation as described in this manuscript.

## Supporting Information

**Figure S1.**
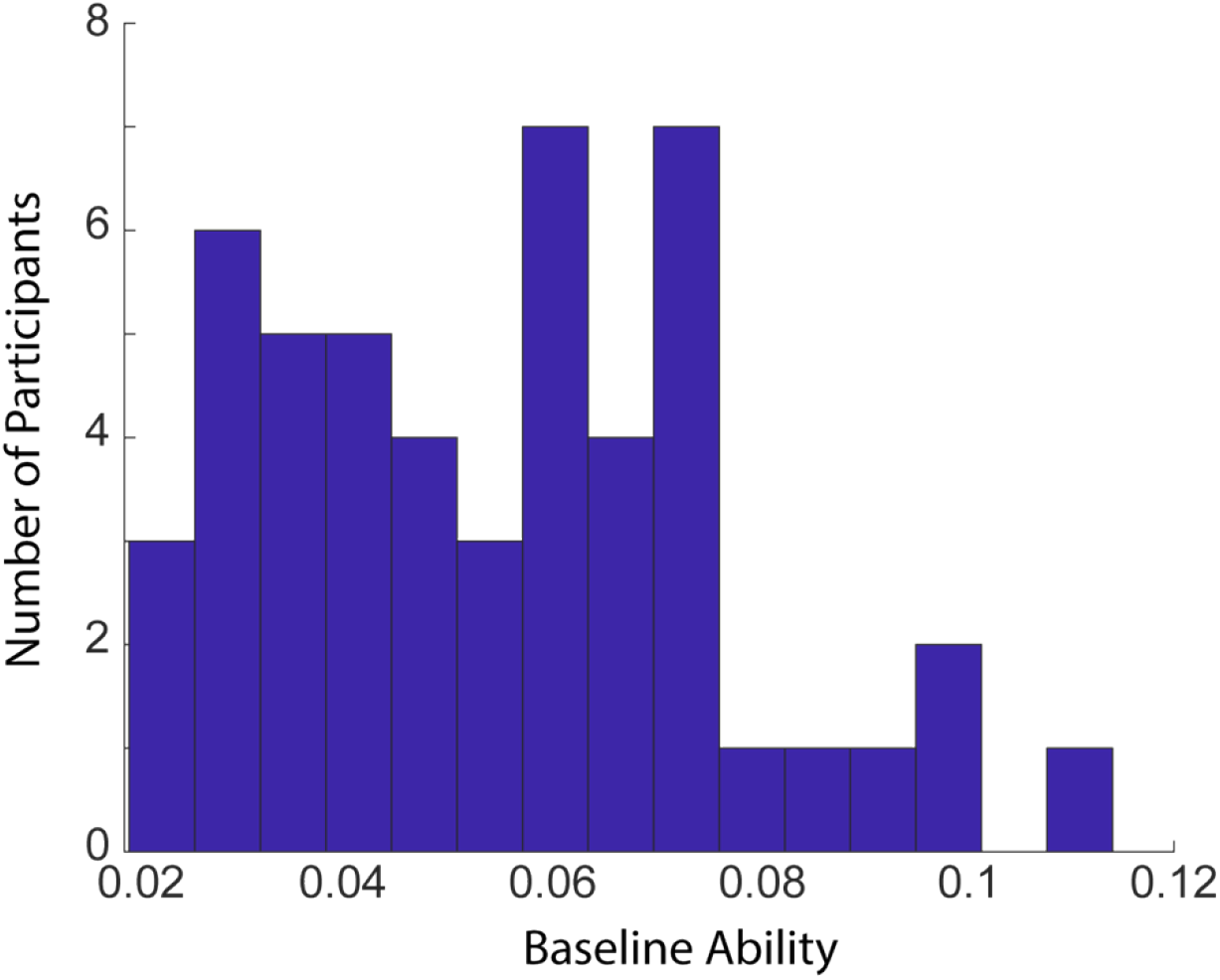
Histogram plot showing the number of subjects in every baseline ability (n = 50). More subjects were on the lower part of the spectrum of the baseline ability range than the higher part.

## Supplementary Results and Discussion

### Sensations and Blinding of tACS

All subjects reported a low level of sensation during stimulation, with no serious adverse side effects, which complies with the previous literature (73). Interestingly, itching and fatigue were more frequently reported than sensations such as burning, pain, phosphenes, and warmth (**Figure S2** in Supporting Information). No reliable correlations were found between the intensity ratings for all sensations and tACS amplitude (all *p* >.10). Moreover, the blinding efficacy of tACS was at chance level (**Figure S3** in Supporting Information). However, the correct indication of real stimulation increased with the current of the applied stimulation.

**Figure S2.**
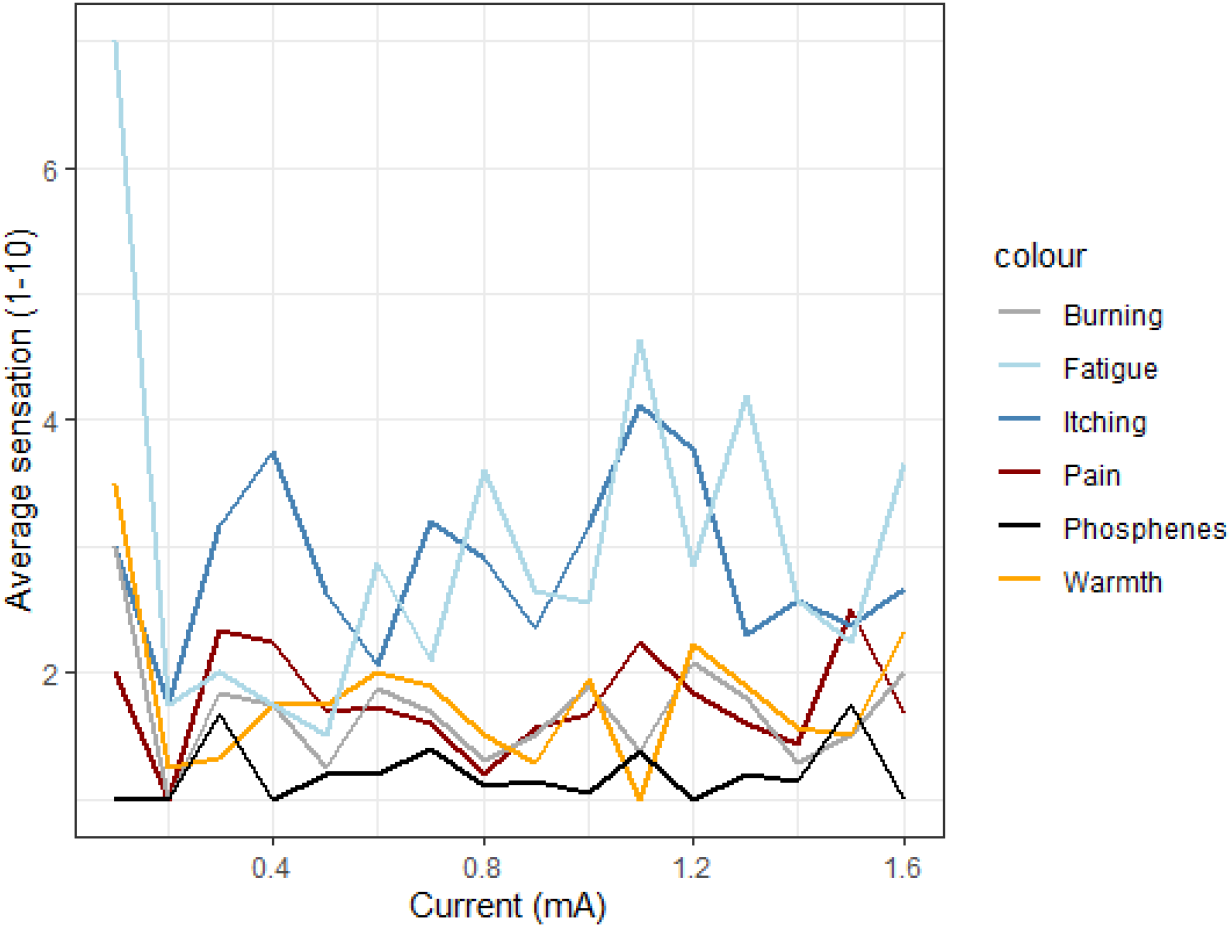
Side effects of different tACS frequency-current combinations. Different side-effects are shown according to the indicated sensation on a scale from 1-10 (n = 150 based on 50 subjects). 1 was indicated as a low sensation (‘I did not feel the sensation’) and 10 is a strong sensation (‘I felt the sensation to a considerable degree’). The high value for fatigue at 0.1 mA is likely to be due to the low number of subjects (n = 2) who received this stimulation, and it might reflect a general state.

**Figure S3.**
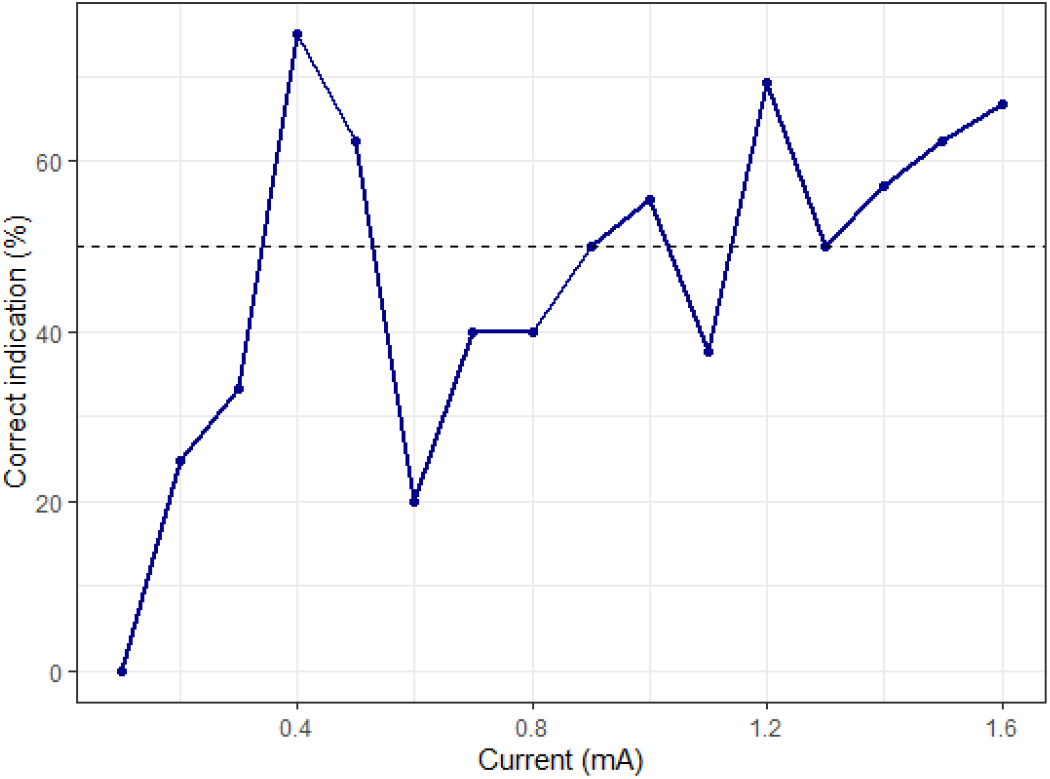
Blinding efficacy of tACS. The figure shows the percentage of correct indications that stimulation was real for every applied current (n = 150 based on 50 subjects). Blinding efficacy of tACS is at change level (∼50%).

**Figure S4.**
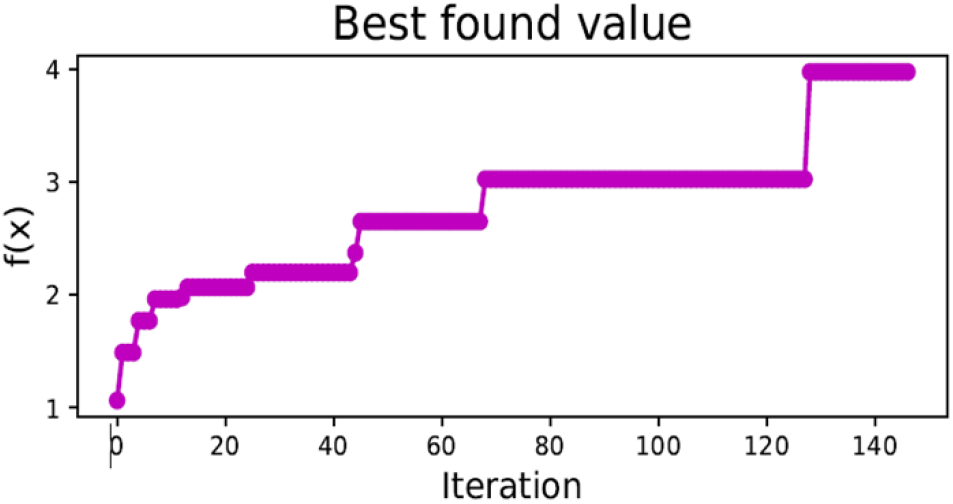
Best found value of the personalized Bayesian optimization (pBO) procedure without exclusion of data point 46 (n = 50). Arithmetic performance in terms of drift rate for every best-found value for *f*(x) and for every iteration of the pBO procedure during stimulation without exclusion of data point number 46.

**Table S1.** Number of subjects (n) according to the different currents (mA) received in one stimulation block.

| **Current**<br>**(mA)** | **N (number**<br>**of subjects)** |
| --- | --- |
| 0 | 0 |
| 0.1 | 2 |
| 0.2 | 4 |
| 0.3 | 6 |
| 0.4 | 4 |
| 0.5 | 16 |
| 0.6 | 15 |
| 0.7 | 10 |
| 0.8 | 10 |
| 0.9 | 14 |
| 1.0 | 18 |
| 1.1 | 8 |
| 1.2 | 13 |
| 1.3 | 10 |
| 1.4 | 7 |
| 1.5 | 8 |
| 1.6 | 3 |

**Table S2.** Number of subjects (n) according to the different frequencies (Hz) received in one stimulation block.

| **Frequencies**<br>**(Hz)** | **N (number**<br>**of subjects)** | **Frequencies**<br>**(Hz)** | **N**<br>**(number**<br>**of**<br>**subjects)** |
| --- | --- | --- | --- |
| 5 | 3 | 29 | 2 |
| 6 | 3 | 30 | 2 |
| 7 | 2 | 31 | 1 |
| 8 | 2 | 32 | 7 |
| 9 | 3 | 33 | 8 |
| 10 | 4 | 34 | 1 |
| 11 | 3 | 35 | 4 |
| 12 | 1 | 36 | 5 |
| 13 | 2 | 37 | 5 |
| 14 | 1 | 38 | 4 |
| 15 | 1 | 39 | 1 |
| 17 | 6 | 40 | 1 |
| 18 | 3 | 41 | 1 |
| 19 | 7 | 42 | 2 |
| 20 | 4 | 43 | 9 |
| 21 | 3 | 44 | 5 |
| 22 | 1 | 45 | 4 |
| 23 | 4 | 46 | 8 |
| 24 | 2 | 47 | 3 |
| 25 | 4 | 48 | 4 |
| 26 | 1 | 49 | 1 |
| 27 | 1 | 50 | 3 |
| 28 | 2 |  |  |

### EEG and Arithmetic Performance

As stated in our pre-registration, we also investigated EEG changes induced by tACS during arithmetic performance (see **Tables S1 and S2** in Supporting Information). To do this reliably, we computed power (electrode Fz) and wPLI scores (electrodes F3 and P3) for each tACS block and normalized these values according to the baseline rs-EEG values. This normalization was undertaken to exclude possible frequency band specific power changes induced by stimulation at a fixed frequency. For example, stimulation at 10 Hz could lead to an increased entrainment in the 10 Hz frequency band (74). A four-way interaction was found between arithmetic baseline ability, EEG power, current, and frequency when predicting arithmetic performance (SE = 0.01, df = 71, t = 3.48, *p* < .001). To look more closely at this interaction, we investigated the interaction between EEG power, current, and frequency in subjects with low and high baseline abilities separately in a mixed effects model by performing a median split (median = - 2.85). We revealed a 3-way interaction for subjects with low baseline ability (n = 25) (**Table S3 and Figure S5** in Supporting Information). In contrast, for high baseline ability subjects (n = 24), there was no three-way interaction present (all *p* > .08) (**Table S4** in Supporting Information). In short, for low ability subjects, a high current (1.6 mA) leads to a steeper increase in arithmetic performance and EEG power when frequencies are low (4 Hz) (**Figure S5** in Supporting Information). This pattern decreased as frequency increased and flipped to a decreased performance and power when the applied tACS frequency was in the 50 Hz frequency range. When running the same analysis for connectivity, no three-way interaction was found for low and high ability subjects (*p* > .05).

Our electrophysiological findings after stimulation indicated an interaction between tACS parameters, oscillatory brain activity, and arithmetic performance for subjects with low baseline ability (**Table S3** in Supporting Information). Interestingly, this interaction between brain stimulation and brain activity was not found for high baseline ability subjects. Our preferred explanation for this finding is that tACS strongly entrains neural oscillations when there is a high-performance gain due to low baseline ability. However, when looking at our group-level pBO model (**Figure 3**) there are no differential effects of tACS on performance levels between subjects with low and high baseline ability. Neural entrainment due to tACS could possibly serve as a compensatory mechanism to improve performance for subjects with low baseline ability (50).

**Table S3.** Fixed effects of the mixed effects model for arithmetic performance in low ability subjects (n = 25). Note: **\*\****p* < 0.05; **\*\****p* <0.01.

| Predictors | Estimates | CI (95%) | df | t-value | p-value |
| --- | --- | --- | --- | --- | --- |
| (Intercept) | 0.04 | -0.43 – 0.52 | 36 | 0.20 | 0.83 |
| EEG power | -0.99 | -1.63 – -0.35 | 36 | -3.13 | 0.003** |
| tACS current | 0.21 | -0.22 – 0.66 | 36 | 1.00 | 0.31 |
| tACS frequency | 0.002 | -0.01 – 0.02 | 36 | 0.36 | 0.71 |
| Power x current | 0.80 | 0.19 – 1.41 | 36 | 2.66 | 0.01* |
| Power x frequency | 0.02 | 0.01 – 0.05 | 36 | 2.83 | 0.007** |
| Current x frequency | -0.003 | -0.02 – 0.01 | 36 | -0.40 | 0.68 |
| Power x current x frequency | -0.02 | -0.04 – -0.00 | 36 | -2.53 | 0.01* |

**Table S4.** Fixed effects of the mixed effects model for arithmetic performance for high ability subjects (n = 24). Note: \*\**p* < 0.05; \*\**p* < 0.01.

| Predictors | Estimates | CI (95%) | df | t-value | p-value |
| --- | --- | --- | --- | --- | --- |
| (Intercept) | 0.05 | -0.37 – 0.47 | 38 | 0.24 | 0.80 |
| Power | 0.53 | -0.08 – 1.16 | 38 | 1.75 | 0.08 |
| tACS current | 0.17 | -0.34 – 0.70 | 38 | 0.68 | 0.49 |
| tACS frequency | 0.004 | -0.01 – 0.02 | 38 | 0.73 | 0.46 |
| Power x current | -0.51 | -1.27 – 0.25 | 38 | -1.36 | 0.17 |
| Power x frequency | -0.01 | -0.03 – 0.00 | 38 | -1.78 | 0.08 |
| Current x frequency | -0.006 | -0.02 – 0.01 | 38 | -0.88 | 0.38 |
| Power x current x frequency | 0.01 | -0.01 – 0.04 | 38 | 1.32 | 0.19 |

**Figure S5.**
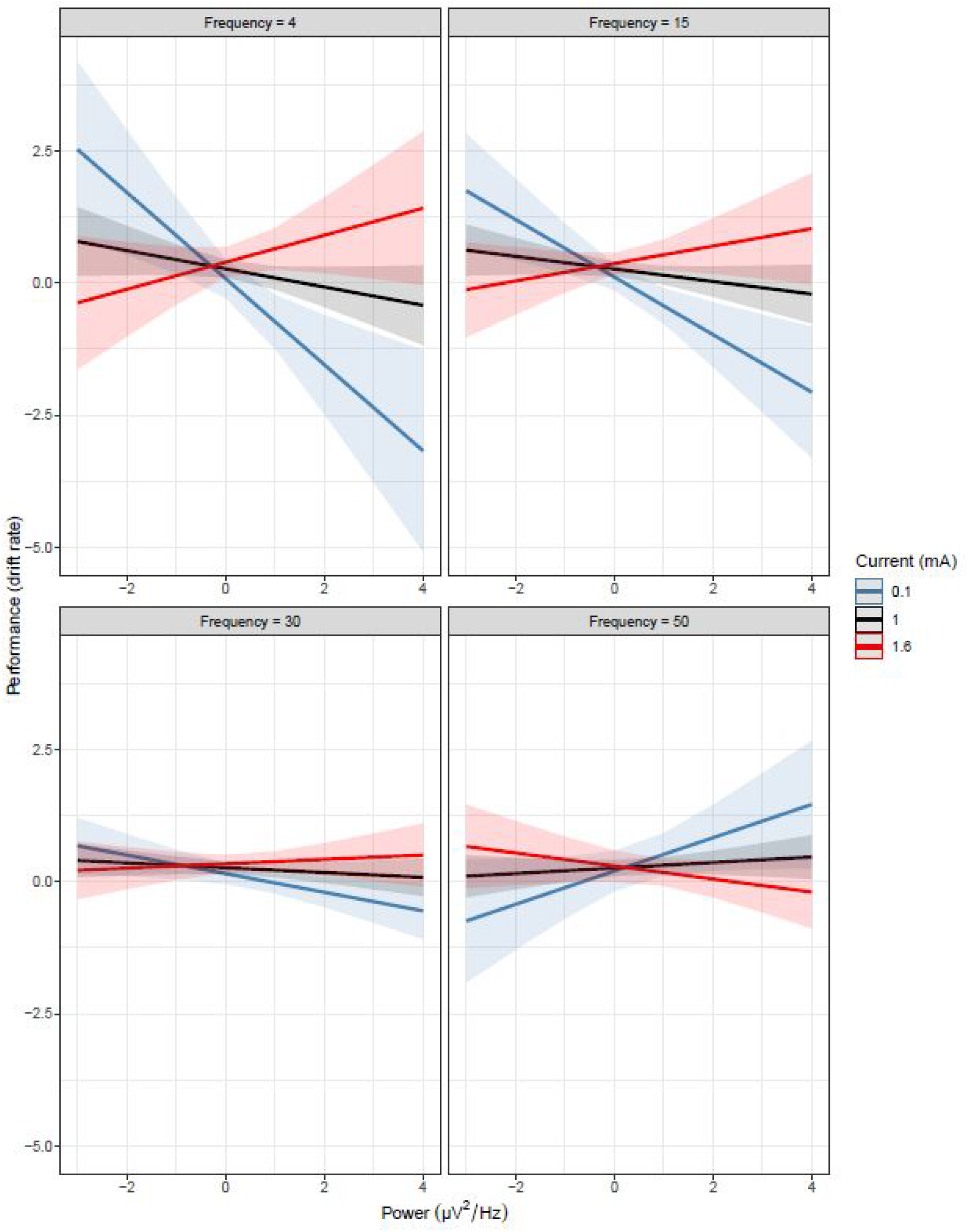
The interaction between EEG power, current, and frequency in predicting arithmetic performance for subjects with low arithmetic baseline ability (n = 25). Arithmetic performance (log transformed drift rates) during stimulation is shown on the y-axis and the normalized (post stimulation/pre stimulation) EEG power μV^2^/Hz (log transformed) based on the applied tACS frequency after stimulation is shown on the x-axis for four different tACS frequencies (4 Hz, 15 Hz, 30 Hz, and 50 Hz). Current intensity is indicated by the blue line (0.1 mA), the black line (1 mA), and the grey line (1.6 mA). Shaded areas indicate 95% confidence intervals. Note that different tACS categories and current intensities are presented for visualization purposes, to allow a better grasp of an interaction that is based on continuous variables.

#### Questionnaire Items

Sensation levels experienced during stimulation

After every block in which the subject received stimulation, the following items were presented:

Do you believe that you received real or placebo stimulation?

1) Real
2) Placebo
3) I do not know
4)

Please indicate whether you experienced any discomfort during the stimulation by typing the corresponding number:

1 = None (I did not feel the sensation)
2-3 = Mild (I mildly felt the sensation)
4-6 = Moderate (I felt the sensation)
7-10 = Strong (I felt the sensation to a considerable degree)

for Pain, Burning, Warmth/Heat, Fatigue/Decreased alertness, Flashing lights

In the case of perceived sensations: How much did these sensations affect your general state?

1 = Not at all
2 = Slightly
3 = Considerably
4 = Much
5 = Very much

How long did the sensations last? (0 to 10 minutes)?

